# Annihilation of action potentials induces electrical coupling between neurons

**DOI:** 10.1101/2023.05.24.542064

**Authors:** Moritz Schlötter, Georg Maret, Christoph J. Kleineidam

**Author notes:** **For correspondence:** (MS).

## Abstract

Neurons generate and propagate electrical pulses called action potentials which annihilate on arrival at the axon terminal. We measure the extracellular electric field generated by propagating and annihilating action potentials and find that on annihilation, action potentials expel a local discharge. The discharge at the axon terminal generates an inhomogeneous electric field that immediately influences target neurons and thus provokes ephaptic coupling. Our measurements are quantitatively verified by a powerful analytical model which reveals excitation and inhibition in target neurons, depending on position and morphology of the source-target arrangement. Our model is in full agreement with experimental findings on ephaptic coupling at the well-studied Basket cell - Purkinje cell synapse. It is able to predict ephaptic coupling for any other synaptic geometry as illustrated by a few examples.

## Introduction

Neurons are characterized by electrically excitable membranes and most neurons in a brain are able to generate action potentials (APs) which are propagating along the axon. At the synaptic terminal they eventually lead to electro-chemical processes, allowing communication with postsynaptic target neurons (***Eccles, 1982***). The most prevalent form of communication between neurons is by transmitter release at the presynaptic terminal, and subsequent binding of these at postsynaptic sites leads to modulation, e.g. excitation or inhibition. Less common in nervous systems of any organism are gap junctions between neurons, resulting in an aggregated functional unit because internal current is transferred from one neuron to the other neuron. Beside these two forms of communication between neurons, the electrical field itself which is generated at an excited membrane can have an impact on the membrane potential of cells in the vicinity (***Jefferys, 1995; Anastassiou and Koch, 2015; Rebollo et al., 2021***). Rapid changes of the membrane potential, as it occurs when APs are generated, induce electrical fields, that by superposition are also present in neighboring cells. The impact of a single, propagating AP on adjacent neurons is very small, nevertheless, along active, parallel fibers synchronization of APs can occur (***Katz and Schmitt, 1940***). When the term ephaptic interaction was coined by ***Arvanitaki (1942***), she concluded that the blockage of APs is the main source of such effects (hereafter: ephaptic coupling) in nervous systems. Indeed, APs arriving at the synaptic terminal of a neuron annihiliate, thereby generating a more potent electrical field for ephaptic coupling. The functional significance of ephaptic coupling at synaptic terminals is well documented in at least two systems: At the Mauthner cell in teleost fish and at the Purkinje fibers of the cerebellum in vertebrates (***Furukawa and Furshpan, 1963; Blot and Barbour, 2014***). In both systems, APs arriving at the synaptic terminal of the source neuron (e.g. the Basket cell of the cerebellum) modulate the initiation of an AP in the target neuron by transiently changing the membrane potential. Although the phenomenon of ephaptic coupling is known since many decades (***Arvanitaki, 1942***), a quantitative physical description is missing. This is even more surprising when considering that ephaptic coupling at synaptic terminals is ubiquitous and may well contribute to communication between many different types of neurons, beside the mentioned Mauthner cell and Purkinje fiber. In our study, we aim to describe the electric field of an AP arriving at the synaptic terminal in such detail that the ephaptic coupling and the change in membrane potential of a known target neuron can be predicted.

The propagating action potential is driven by a voltage dependent transition of the membrane from a resting to an excited state. As a consequence of electro-neutrality, current flows across the membrane resulting in a closed loop through extracellular and intracellular medium. The current loop includes the resistive extracellular medium where the AP generates a voltage signature. In 1952, Hodgkin-Huxley introduced a model (HH model) for generation and propagation of APs (***Hodgkin and Huxley, 1952***). It is based on the cable equation and incorporates Nernst equations for different ions and changing ion specific conductances upon activation of ion channels. The HH model is a powerful tool to study and evaluate the properties of ion channels in excitable membranes on a microscopic level, and the great attention it received since being introduced is more than justified (***Catterall et al., 2012***). However, the macroscopic phenomenon of current loops and associated electric fields is already well accounted for by a much simpler model introduced by Tasaki and Matsumoto (***Tasaki and Matsumoto, 2002; Tasaki, 2006***) (TM model) without the need of a multitude of specific parameters as required for example in the HH model. While the TM model can be considered as a simplification of the HH model, the mechanism that the authors had in mind is very different (***Tasaki, 2002; Tasaki et al., 1965, 1971***). In general terms, neural membranes can go through transitions where the electric properties change extremely fast (***Fedosejevs and Schneider, 2022; Mussel et al., 2021; Horkay et al., 2000***). The TM model describes a resting and an excited state, each with a linear cable model. It uses the cable equation with two state-specific parameters, its equilibrium potential and conductivity, which undergo a sudden (voltage dependent) switch at the boundary between resting and excited state. The boundary between an excited and a resting section propagates towards the resting side. The AP in the TM model has only 3 degrees of freedom, e.g. its propagation velocity, its amplitude and its length. Nevertheless, the TM model is well suited to describe the current distribution that is driven by the inhomogeneity of local electric fields in the internal and the external medium. Thus, the TM model possibly already is sufficient to describe the mutual influence of neurons by ephaptic coupling, using only three parameters for each neuron.

It is therefore of prime importance to evaluate the quantitative description of the TM model based on experimentally derived values. We explore the behaviour of APs at boundaries to assess these three parameters. A suitable boundary condition (intracellular, axial current equals zero) can be generated experimentally by a collision of two counter-propagating APs (***Tasaki, 1949; Spach Madison S***. ***et al., 1971; Shrivastava et al., 2018; Shrivastava, 2018***). Within any cable model, the very same boundary condition also exists within the axon at the synaptic terminal due to the broken translation symmetry for the current loops (***Spach and Kootsey, 1985***).

In this study, we first focus on collisions of APs. Our experimental observation of colliding APs provides unique access to the spatial profile of the extracellular potential around APs that are blocked by collisions and thus annihilate. Our procedure is as follows: Recording propagating APs allows to determine both the propagation velocity and the amplitude of the extracellular electric potentials. The collision experiment provides additional information on the characteristic length *λ*^⋆^, thereby fully determining the parameters of the TM model. We find that TM is sufficient to describe the onset of the electric field dynamics of an AP. However, as TM does not include the slow return of the excited membrane to the resting state we introduce an extension of the TM model that we call the Relaxing Tasaki Model (RTM) which adds an empirical slow relaxation term to the TM model. We find that RTM accounts for the full spatio-temporal signature of propagating and colliding APs and is therefore also very useful to predict electric effects at axon terminals e.g. at synapses. Since the RTM model sets a complete framework to describe APs and to predict ephaptic interactions for a given morphology, the RTM model is used in the last part of this paper to simulate various representative pre- and postsynaptic morphologies and geometries. Around AP propagation boundaries, we find excitatory as well as inhibitory regions, depending on timing, relative position, orientation and morphology of source and target neuron. We expect these predictions to be robust, because the model is absolutely minimal and matches the experimental observations.

## Results

### Extracellular potential around AP collisions

We record the extracellular electric potential of APs, generated by the median giant fibers (MGF) of the ventral nerve cord of an earthworm using the setup shown in ***Figure 1***A and B. In a first experiment the ends are stimulated individually and the APs are detected at several positions along the fiber. We verify the vitality of a nerve cord by asserting that both fibers (MGF and the lateral giant fibers (LGF)) reliably conduct APs in both directions, and in our experiments, we investigate selectively the MGF only. The signatures ***V***_*e*_(*t*) of both orthodromic and antidromic propagating APs look indeed very similar. We observe a biphasic voltage spike propagating with 14.5(11) m*/*s (***Figure 1***A).

**Figure 1.**
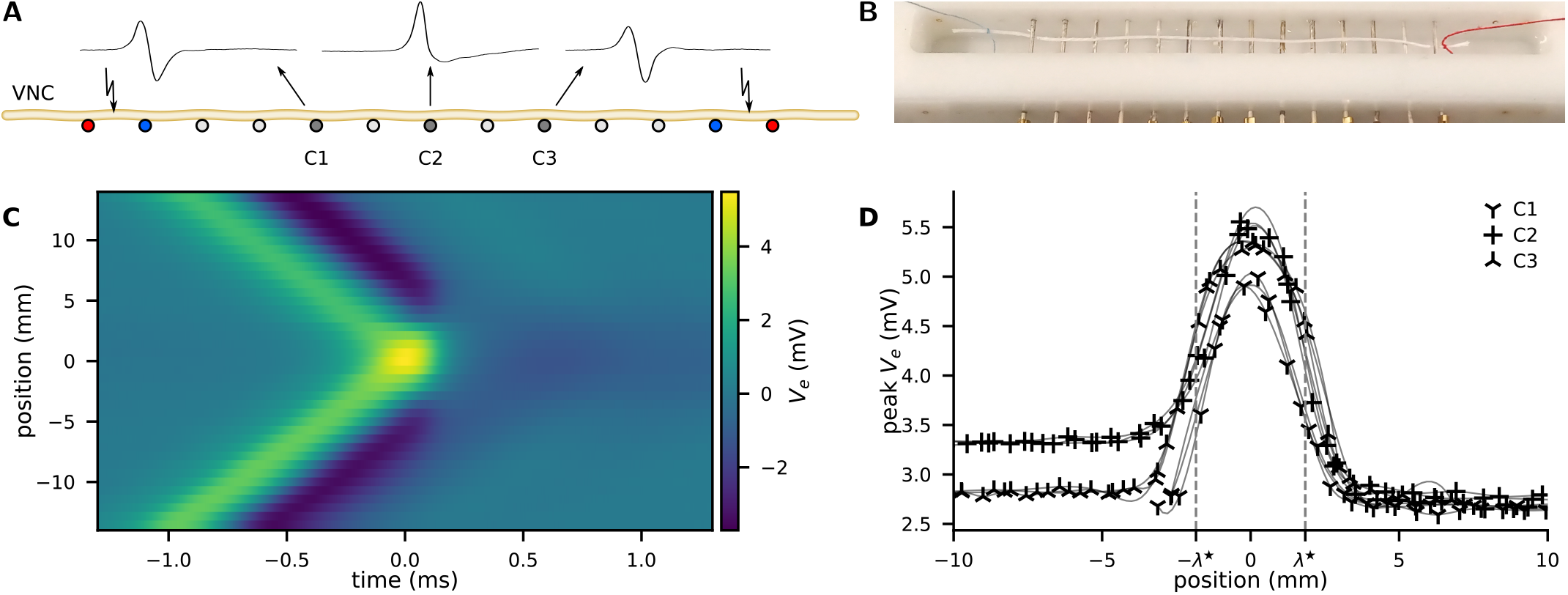
AP collision experiment. (A) The ventral nerve cord (VNC) of an earthworm is positioned on a series of transverse, single-ended electrodes to excite and monitor APs. Individual or two counter-propagating APs are generated by stimulating the nerve at the outermost electrodes. Some of the inner electrodes (C1, C2, C3) are used for recording, all other electrodes are grounded. Propagating APs generate biphasic electrical potential as sketched above C1 and C3 whereas colliding APs provoke an essentially unipolar peak (cf. C2). By variation of the delay between the opposing stimuli, the collision can be generated anywhere along the nerve between the stimulation electrodes. (B) Photograph of the recording chamber with a white thread to illustrate the position of the VNC. (C) Extracellular recording of propagating and colliding APs. A collision sweep experiment yields multiple recordings with varying distance between recording site and point of collision. The collision is captured in the recording line at y-position 0 mm, while orthodromic propagation is at the top and antidromic propagation is at the bottom. (D) The peak amplitude as a function of the distance to the collision. Examples of four sweeps at three positions along the nerve cord. As a guide to the eye, the data points are connected by a cubic spline (thin lines). Figure 1—figure supplement 1. Raw Data Figure 1—figure supplement 2. Fit of the collision width

In the main experiment, both ends are stimulated simultaneously and the APs collide close to the central recording electrode. We observe that the APs do not penetrate each other but always annihilate at the collision site (***Figure 1***A) (***Tasaki, 1949***). At this position, the trace of ***V***_*e*_(*t*) becomes more monophasic, with an almost doubled positive peak amplitude (***Figure 1***A). The negative phase of ***V***_*e*_(*t*) at the collision site is considerably diminished and shows a distinct slow relaxation. The peak amplitude of propagating APs is 2.9(2) mV while the colliding APs peak is 5.2(3) mV.

The width of the collision is a measure of the characteristic length *λ*^⋆^ of the AP and is uniquely revealed by a collision sweep experiment. We experimentally positioned the site of collision along the neuron by introducing a delay between the opposing stimuli. Assuming symmetric propagation velocities, a delay Δ*t* displaces the collision by *x* = *v*_*p*_Δ*t/*2 where *v*_*p*_ is the propagation velocity and *x* is the distance between the collision and the recording electrode. The spatial extend of the collision process is then found by repeating the experiment at various delays Δ*t* between the opposing stimuli, see (***Figure 1***C). The propagation velocity is derived from the time of arrival at separate recording sites. The peak amplification can be used as a measure of the width of the collision (***Figure 1***D). We use four recordings at three positions (anterior, medial and posterior) along the nerve cord and we assess the mean full width at half maximum as 3.8(5) mm.

### Model of the action potential

We describe a neuron by a classical cable model as shown in ***Figure 2***A (see “Methods” for further details). Such a model consists of a chain of RC circuits composed of resistors with conductivity *g*_*m*_ (S*/*m) and capacitors *c*_*m*_(F*/*m) which are connected by an inner resistivity *r*_*i*_ (Ω*/*m). The inner axial current ***I***_*ax*_ is driven by the gradient of the potential ***V*** (*x, t*) inside the neuron. A common approach is to assume a constant capacity and to neglect external fields and inhomogeneities in the neuron. Then, the balance of currents results in the cable equation with the membrane potential ***V***_*m*_

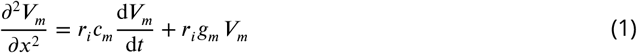

**Figure 2.**
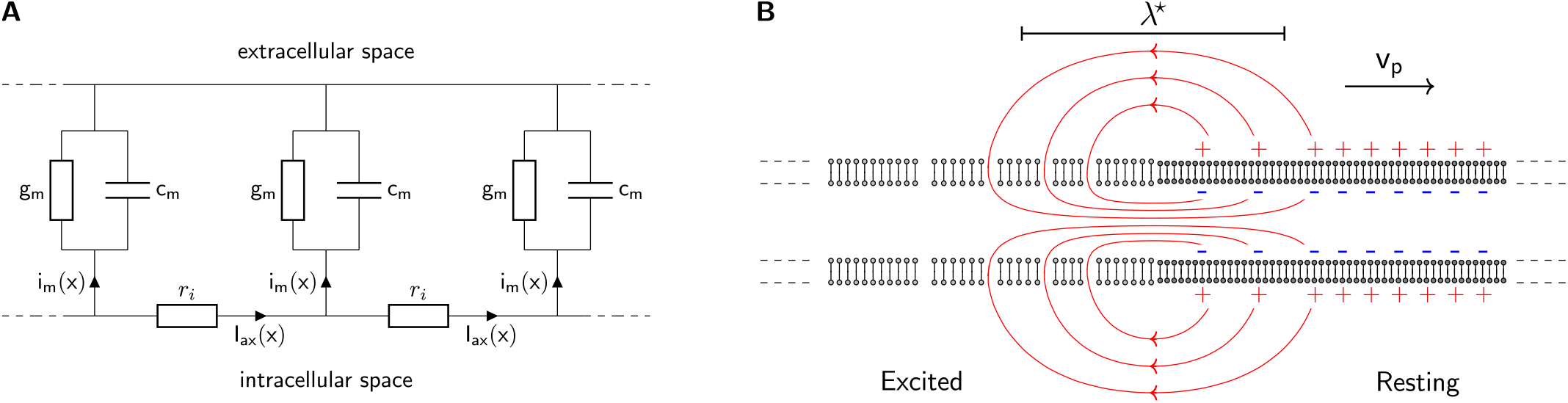
The classical cable model. (A) A chain of identical RC circuits with conductivity *g*_*m*_, capacity *c*_*m*_ and inner resistivity *r*_*i*_ mimics the electric properties of the cellular membrane and connects the extracellular and intracellular space. The change of inner axial current ***I***_*ax*_(*x*) is given by the transmembrane current density *i*_*m*_(*x*) (Kirchhoff’s current law). (B) The propagating solution of the Tasaki-Matsumoto model. A boundary between a resting state and an excited state induces an axial current, that causes a propagation of the boundary. The axial current flows in closed loops and returns within the resistive extracellular medium, causing an extracellular potential that travels with the current source along the neuron.

The first term on the right describes the capacitive charging by an axial current which is linked to the rate of change in membrane potential. The second term describes the contribution of resistive membrane current to axial current. The relationship between the axial current ***I***_*ax*_ and the membrane current *i*_*m*_ is further described in the Methods and Materials eq. 5.

APs are characterized by a solitary i.e. non-spreading spatio-temporal shape. In the cable model, the generation of such APs requires an additional non-linear response of the membrane. Commonly, living cells are electrically charged to a negative equilibrium potential ***V***_*eq*_ across their membrane. In excitable cells, e.g. neurons, a change of the membrane potential beyond a critical threshold value leads to a rapid transition from the resting into an excited state of the membrane which essentially differ in their respective resistivities. The excited state, with drastically increased conductivity, is indicated by the starred values 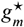 and potential 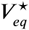.

A local transition of a membrane from the resting to excited state results in a boundary between ***V***_*eq*_ and 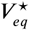. The difference in membrane potentials between the resting and excited state induces an internal axial current. The increased conductivity at the excited state *drives* the internal current and depolarizes the adjacent membrane. The flow lines of internal, transmembrane and external current form a closed loop around the position of the transition as sketched in ***Figure 2***B. All internal current is present as an extracellular return current and in neurons, this process produces a propagating, dipolar local current source.

The Tasaki-Matsumoto (TM) model describes both the resting and the active state by linear cable models (***Tasaki and Matsumoto, 2002; Tasaki, 2006***). The resting state has a negative electrochemical equilibrium potential ***V***_*eq*_ = −100 mV and a very low conductivity *g*_*m*_. In the excited state the conductivity is drastically increased to 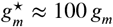 and the equilibrium potential 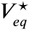 is zero. The transition from the resting to the excited state is discontinuous and occurs at ***V***_*m*_ = −50 mV. The TM model does not address how the resting state is maintained before a transition, nor does it address how the resting state is restored thereafter.

A simple analytical solution of the TM model describes the propagating initial depolarization of a nerve signal. This includes an exact expression for the propagation velocity *v*_*p*_, and for the spatial length *λ*^⋆^ of the initial current dipole as illustrated in ***Figure 2***B. The velocity is determined by the delay in charging of neighboring capacitors, while the leak conductivity barely effects the process. Therefore, we base our study on simplified expressions by neglecting the leak conductivity, as it amounts to less than 1% of the conductivity in the excited state (***Tasaki and Matsumoto, 2002***). Then the analytical solution of the TM model yields

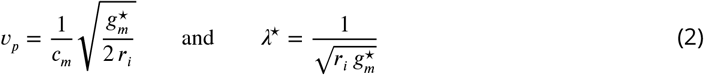

Note that the length scale *λ*^∗^ used here is not equivalent to *λ* used in basically all previous models where *λ* denotes the damping length due ionic leakage neglected here. The products in eq. 1 can be expressed as *r*_*i*_ *c*_*m*_= (*v*_*p*_ *λ*^⋆^)^−1^ and 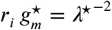. According to eq. 1 (*r*_*i*_ *c* _*m*_) ^−1^ is the diffusion constant of the spreading potential ***V***_*m*_ which connects a typical time scale *τ*_*L*_ with a typical diffusion length *L* by *τ*_*L*_ = *r*_*i*_ *c*_*m*_***L***^2^. The two products *r*_*i*_ *c*_*m*_ and 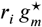 are directly determined in our experiment from measurements of *v*_*p*_ and *λ*^⋆^, and for a comparison of our AP measurements with the TM model only one degree of freedom is missing, which is the amplitude of the AP. At this point wefit the amplitude of the model to our experimental recording by estimating an appropriate extracellular conductivity.

The fast switch from resting to excited state in the generation of an AP is followed by a slow process of repolarization that restores the resting state. In order to provide a more general model that also accounts for repolarization we add a simple extension to the TM model and refer to this as Relaxing Tasaki Model (RTM). We approximate the repolarization in the extracellular potential by extending the TM model with an (ad-hoc) exponential repolarization function (see “Methods”).

### Comparison of measurements with model simulations

The above collision sweep experiment provides a detailed survey of the extracellular electric signature of APs. We now compare these results to the TM model, the RTM model and the classical HH model (see “Methods”) by simulating a nerve of length 10 cm with 2000 compartments (Conductivities in the HH model are area specific, here we use a diameter of 80 µm). All models are adjusted to the MGF with identical procedure. The parameters *r*_*i*_ and *c*_*m*_ are used to match the propagation velocity *v*_*p*_ and the width *λ*^⋆^ of the collision to the experimentally determined values. Thefitting procedure is as follows: (i) Propagating APs are simulated. For a given *r*_*i*_ the velocity is adjusted using *c*_*m*_. (ii) An AP collision is simulated and the width is compared to the experiment (see ***Figure 1—figure Supplement 2***). If the width deviation is below 1%, the model is accepted. Otherwise *r*_*i*_ is updated and the procedure is repeated from step (i). The TM model obeys the analytical expressions and converges immediately. The RTM model deviates from the TM expressions only by a few percent and converges very fast. The HH model takes more iterations to converge. In_192_ all cases, the procedure is unambiguous and stable. The complete parameters for all models are given in ***Appendix 3 — Table 2***.

While the individual values of *r*_*i*_ and *c*_*m*_ depend on the choice of *g*^⋆^, the product *r*_*i*_*c*_*m*_ does not. From our measurements we obtain *r*_*i*_*c*_*m*_ = 26 s*/*m^2^, which is in good agreement with literature values (***Tasaki and Matsumoto, 2002; Tasaki, 2012***). The resulting length parameter is *λ*^⋆^ = 1.8 mm which is about half the width of the peak in ***Figure 1***D. The deviations between the TM model and the experiment are: 1. Propagating APs cause a negative peak greater than the positive peak (***Figure 3***A). 2. There is a small but sustained negative phase at the collision site (***Figure 3***B). The repolarization time in the RTM model is found by comparing the model to the experimental traces of the colliding AP (***Figure 3***B). A value of *τ*_*r*_ = 0.5 ms provides an acceptable fit. All models allow the reconstruction of the extracellular field of propagating APs within satisfying accuracy.

**Figure 3.**
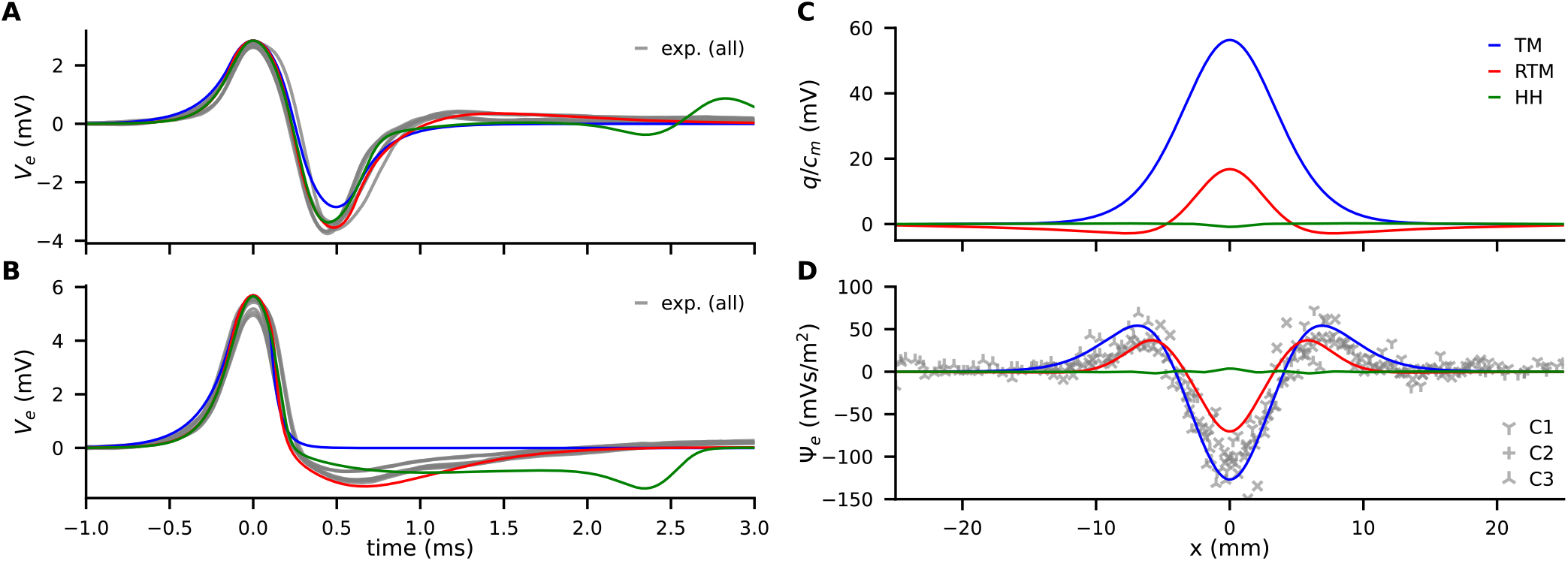
Comparison of experimental data (gray) with models (color, see legend). (A) Traces of ***V***_*e*_(*t*) from propagating APs. (B) Traces of ***V***_*e*_(*t*) at the collision site. (C) Model predictions of the discharge (*q*(*x*)) generated by AP annihilation and released around the collision site. (D) The extracellular discharge of annihilating APs, measured and calculated. The gray crosses are the experimental values from 4 recordings at 3 separate sites along the MGF, where each recording consists of up to 46 traces with varying delay times.

### Discharge of colliding APs

In an active axon, current always flows in and out in equal amounts (electroneutrality) (***Barbour, 2020***). As long as APs propagate the current must sum up to zero in space at any time and in time at any position. However, AP propagation is blocked when the axial current ***I***_*ax*_ is shut down at a boundary condition, e.g. by reaching the axon terminal or by AP collision. Such annihilating APs generate an effective net charge expelled into the extracellular space because the local sum in time can deviate from zero while the spatial sum remains zero. The associated current pulse may act like a stimulating electrode and as current source for ephaptic coupling. In order to access how much charge is expelled, we can calculate the charge as the integral of membrane current over time.

The membrane current *i*_*m*_ is given by the cable equation (eq. 1) as the curvature of the membrane potential 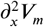, devided by *r*_*i*_. Thus, the discharge *q*(*x*) is the total charge expelled at a boundary (site of collision or axon terminal) and described by

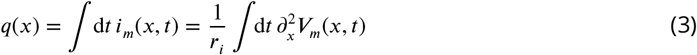

The discharge *q*(*x*) can be calculated with our models and yield very different predictions about its magnitude (***Figure 3***C). It has to be noted that *q*(*x*) does not directly predict ephaptic coupling.

However, *q*(*x*) generates a curvature in the extracellular potential and in order to quantify possible impact of current, we integrate the curvature in the extracellular potential over time and define (**Ψ**) as

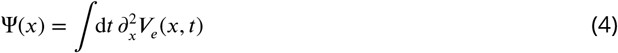

***V***_*e*_ and thus **Ψ** can be assess with our collision measurements and can be used for model validation by comparing the models’ predictions. In our simulations with a multi compartment model, **Ψ** is calculated when APs are colliding, based on parameterization from propagation velocity (*v*_*p*_) and width of the collision (length *λ*^⋆^, see above). ***Figure 3***D illustrates that the TM model is in excellent agreement with the experiment and the RTM model only slightly underestimates **Ψ**. From this observation we conclude that the discharge is driven by the rapid onset of the AP and that the contribution by the repolarizing phase is small. On the other hand, the HH model largely disagrees with the experiments both in the magnitude of the discharge and in its temporal shape.

## Model of ephaptic coupling

First, we briefly investigate the effects of propagating APs on neighboring parallel cells which may give rise to synchronization (***Katz and Schmitt, 1940; Kriebel et al., 1969; Goldwyn and Rinzel, 2016***). Then, we focus on synaptic terminals i.e. situations where the (finite) electric discharge of *annihi-lating* APs may influence surrounding target neurons. We showed above that colliding APs emit a charge *q*(*x*) and can act like a stimulating electrode. Within any cable model, the same boundary condition with ***I***_*ax*_ = 0 also exist at axon terminals (***Plonsey, 1977; Spach and Kootsey, 1985; Kléber and Rudy, 2004***). It was our original idea to conduct the collision experiments and apply the multi compartment simulations to then transfer the knowledge to annihilating APs at axon terminals. Likewise to our simulations for compartments around the site of collision, we can calculate the extracellular potential around the last compartments at the axon terminal. Knowledge of the generated inhomogeneity of ***V***_*e*_ allows to calculate the response of a target neuron that may be close to a site of AP annihilation. The details of this calculation are provided in the methods section. Briefly, the above defined discharge from annihilation at terminals or collisions generates a distortion of the extracellular potential ***V***_*e*_ which in turn induces current in nearby target neurons.

In ***Figure 4*** we show simulations of ephaptic coupling in three different geometries: parallel neurons, end-shaft synapses and end-end synapses. We use common literature values for the inner resistivity *ρ*_*i*_ = 1 Ω m and membrane capacity *c*_*m*_ = 10 mF*/*m^2^ (***Cole, 1975; Carpenter et al., 1975; Bekkers, 2013***). Estimates for the extracellular resistivity are rare and highly variable, in the range of 3 Ω m to 600 Ω m (***Weiss et al., 2008; Lindén et al., 2011***). We derive ρ_*e*_ = 100 Ω m by an estimate based on ***Blot and Barbour (2014***), as explained in ***Appendix 1***

**Figure 4.**
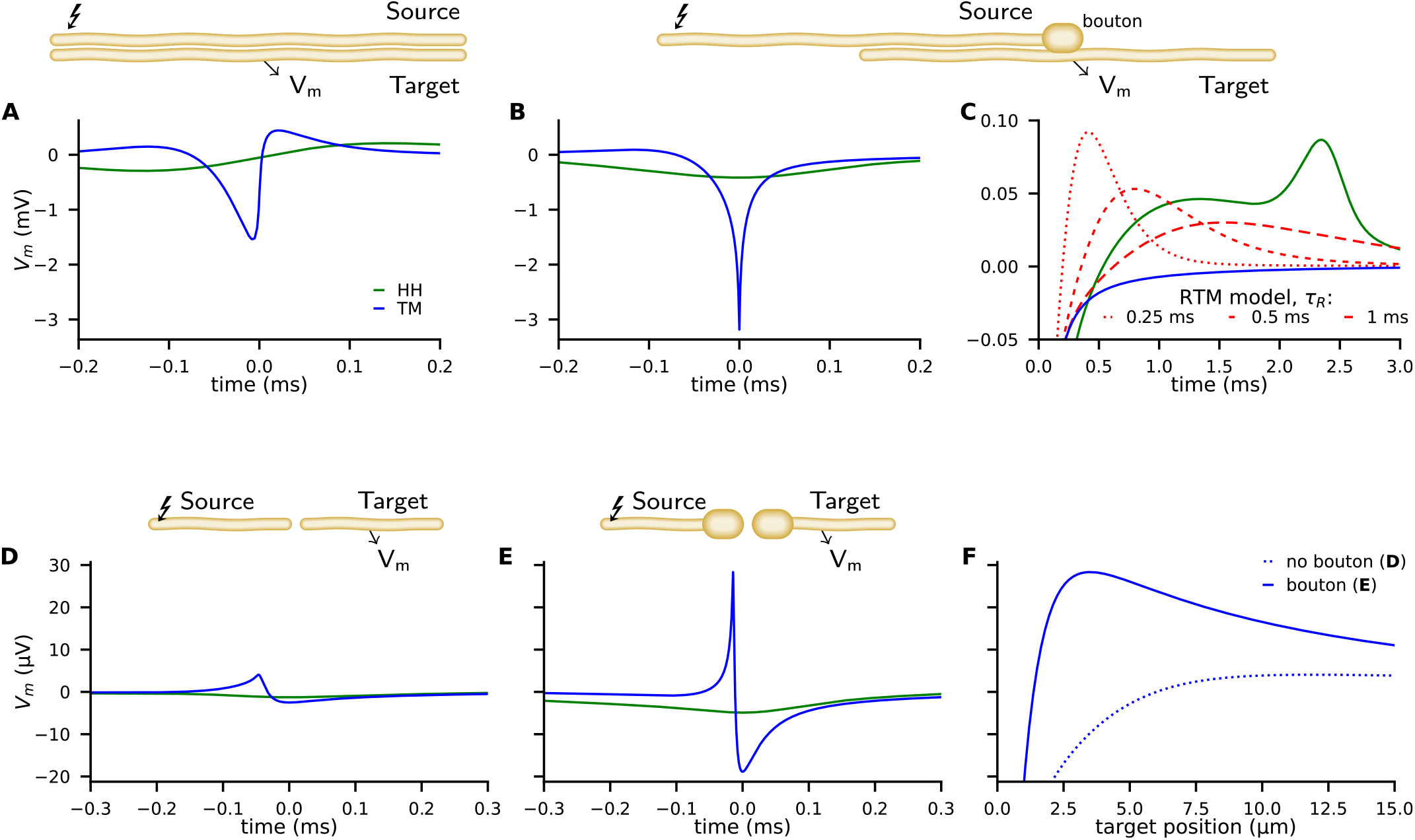
Examples of ephaptic coupling, calculated with the TM-model (blue) and the HH-model (green): (A) in a parallel target neuron when an AP is propagating in the source. (B,C) when the AP is annihilating at a bouton of a neuron terminal (upper neuron in end-to-shaft geometry, similar to the Basket cell – Purkinje cell synapse). The source and target neurons are 1 µm in diameter and are separated by 1 µm (2000 compartments each, length 1 mm, bouton size 2 µm. The neurons are placed next to each other, that is to say the numeric point compartments are separated by 1 µm. Traces denote the target membrane potential next to the point of axon termination. The initial hyperpolarization effect may be followed by a subsequent depolarization, depicted by the RTM model (red) for different relaxation times. (D-F) Ephaptic coupling in an end-to-end synapse, illustrating the enhanced ephaptic coupling, due to enlarged neuron terminals (boutons). Here the source and target neurons are 100 nm in diameter (2000 compartments each, length 300 µm, bouton size 400 nm). The target neurons are 1/4 in length and N (500 compartments, length 75 µm). The TM model generates a distant depolarization. Traces in D and E show the membrane potential of the target at the point where the TM model provokes its maximal depolarization (corresponding to the peaks in F). Traces in F show the spatial profile of the membrane potential along the target, at the time of maximal depolarization (corresponding to the peaks in D, E). Figure 4—figure supplement 1. More details of the end-shaft synapse simulation Figure 4—figure supplement 2. More details of the end-end synapse simulation

The peak membrane conductivity in the TM and RTM models is *g*^⋆^ = 450 S*/*m^2^, and the HH model is used with all classical parameters as specified in the supplementary material (e.g. 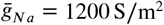). ***Figure 4***A shows the potential ***V***_*m*_(*t*) of the target neuron at the position marked by an arrow when excitation of a propagating AP occurs in the source neuron above. While the response in the target neuron calculated by the HH model is weak and broad, it is more asymmetric, peaked, shorter and stronger in the TM model. Next, we consider an axon terminal that ends near to a neighboring, parallel target fiber. The chosen geometry as sketched in ***Figure 4***B is a coarse representation of the synapse between Basket cell and Purkinje cell with the synapse embedded in a highly resistive structure, called the Pinceau (or basket) at the axon initial segment (AIS). The TM and RTM model predict a strong and sharp initial hyperpolarization of the target neuron while the HH model predicts a comparably weak and smooth effect (***Figure 4***B, see also ***Figure 4—figure Supplement 1***). In the RTM and HH models, this initial effect is followed by a more or less pronounced slow depolarization of the target neuron (***Figure 4***C). The RTM model also reveals that a faster repolarization of the source AP increases the late depolarization of the target neuron.

The modulation of activity in Purkinje cells, e.g. changes in the rate of action potentials in response to an AP arriving at the Pinceau of the Basket cell was described in detail by ***Blot and Barbour (2014***). Since the relationship of membrane potential and the rate of APs in Purkinje cells is reported by the authors to be around 100 Hz*/*mV we can compare our model predictions to the measured rate-modulation of APs. We mapped the result of the models’ predictions for different morphologies to the experimental data (***Blot and Barbour, 2014***) of AP-rate modulation in Purkinje cells (***Figure 5***). Similarly to ***Blot and Barbour (2014***), we used a gaussian filter with 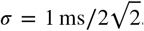. In our two models (TM and RTM), the modulation of not terminating but propagating APs along the source axon on the AP rate of the target cell is minute (***Figure 5***A). Note that this geometry does not correspond to the Purkinje cell - Basket cell connectivity. For annihilating APs at the axon terminal, with excitable segments up to the very end, our models reveal a moderate modulation, and only about half of what was reported for the Purkinje cell by Blot and Barbour (2014). This illustrates the importance of AP annihilation for ephaptic coupling (***Figure 5***B). The Basket cell axon branches, forming the basket and the branchlets or Pinceau embrace the initial part of the Purkinje cell axon. At the Pinceau a different set of ion channels is expressed and it is considered as being non-excitable (***Laube et al., 1996; Southan and Robertson, 2000***). We implemented the morphology of the Pinceau (increased volume and surface area) by implementing a bouton (diameter: axon 1 µm, bouton 2 µm) and non-excitable terminal segments of 15 µm total length (***Bobik et al., 2004***). AP propagation is terminated before reaching the Pinceau terminal and the resulting modulation of AP rate in Purkinje cells highly depends on the morphology of the Basket cell and on *τ* of the RTM model. Non-excitable segments at the last 15 µm result in very consistent predictions from both of our models (TM and RTM) for the reported inhibition and even the rebound activity (between 0.5 and 1.5 ms) is very well predicted by the RTM model, using a *τ* of 0.5 ms (***Figure 5***C). Beside the end-to-shaft geometry as in Basket cell – Purkinje cell connections, other morphologies and geometries are common in neural networks. We consider target neurons pointing towards the site of annihilation (***Figure 4*** D-F, see also ***Figure 4—figure Supplement 2***). The geometry is reminiscent of an excitatory synapse, where an axon terminal is facing towards a dendritic spine. We choose the same parameters that we used for the end-shaft synapse, but the target neuron is placed in projection of the source nerve with a gap in between them of 10 nm. We include enlargements at the ends since boutons are common at the source and target neuron. Such enlargements significantly amplify ephaptic coupling. We observe that the strong hyperpolarization of nearby postsynaptic membranes is accompanied by a distant depolarization. In ***Figure 4*** D,E we present the time course of the induced membrane potential in the target at the point of maximal depolarization. In ***Figure 4*** F the spatial profile of the induced voltage is given at different times.

**Figure 5.**
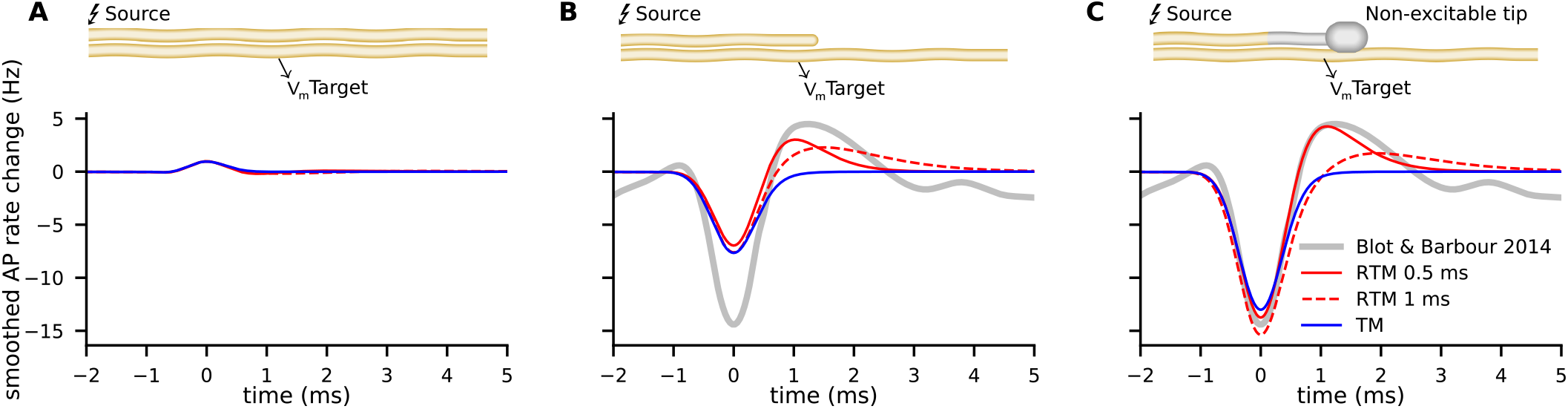
Comparison of experimental data from ***Blot and Barbour (2014***) on modulation of Purkinje cell activity (grey lines) with our RTM model predictions (red lines with two different *τ*) and the TM model prediction (blue) for three different geometries and physiological properties of the source neuron (e.g. the Basket cell). (A) excluding the annihilation of an AP at the source neuron, i.e. a propagating AP, cause only minute modulation of the predicted AP rate in the target neuron. Note that this example does not represent the Basket cell terminal with annihilating APs. (B) annihilation of an AP at the terminal of the source neuron, with all segments being excitable in our calculation, cause moderate modulation. (C) source neuron with similar properties to the Basket cell, i.e. a bouton and last segments non-excitable (corresponding to 15 µm with no switch from resting state to excited state), cause inhibition and rebound that is very similar as described by ***Blot and Barbour (2014***).

The total amplitude of ephaptic coupling strongly depends on the choice of parameters. If we use *g*^⋆^ = 1200 S*/*m^2^ (as commonly used in the HH model) instead of the 450 S*/*m^2^ used in ***Figure 4***, the effect more than doubles. Note that much higher values up to 30 kS*/*m^2^ have been reported for specific locations (***Holt and Koch, 1999***) which suggests even stronger ephaptic interactions in certain situations. The distant depolarization shown for the end-end synapse (***Figure 4*** D,E) is only a few tens µV, but as the amplitude of synchronized input from many dendrites adds up and cerebellar neurons have thousands of dendrites we may expect depolarisation signal strengths in the mV range.

## Discussion

Our experimental design of a custom made recording chamber with a nerve, hanging over a series of electrodes allows accurate measurements of the space and time dependent extracellularfield ***V***_*e*_. Since, according to eqs. 9, variations of ***V***_*e*_ mirror variations of the membrane current *i*_*m*_, we obtain information on *i*_*m*_, on the associated velocity *v*_*p*_ and length scale *λ*^⋆^ of APs. Both of them are measured with great accuracy (about 10%), which allows us critical bench-marking of current theoretical models of AP’s. We find that both, the Tasaki-Matsumoto model and the Hodgkin-Huxley model are well suited to describe the extracellular electric potential of propagating APs which have symmetric forward and backward flow of *i*_*m*_.

However, when an AP encounters a boundary, either at a axon terminal or in a collision with a counter-propagating AP, the current flow and ***V***_*e*_ become asymmetric. In fact, AP propagation is terminated when the overall local axial current is annihilated by a collision event and at axon terminals. At this point, the lack of internal stimulating current causes APs to disappear, as already described by ***Tasaki (1949***). During AP annihilation, the extracellular potential becomes a distinct monophasic spike. This phenomenon has been observed for artificially produced as well as naturally occurring collisions in various systems (***Spach Madison S***. ***et al., 1971; Steinhaus et al., 1985; Tasaki, 1949, 1955***).

We describe the annihilation process by the time integrated charge (*q*, eq. 3) which is expelled at boundary condition, e.g. the axon terminal. From our collision experiments, we extract the spatial shape of the generated extracellular potential. The impact of extracellular potentials that are applied with microelectrodes often is described with the term ‘activating function’ (***Rattay, 2008***). We extended this view by the integration of the naturally occurring ‘activating function’ of neurons over time (**Ψ**, eq. 4).

We find that the experimentally observed values of **Ψ** and thus the underlying discharge *q* can be predicted very well by the TM/RTM models, while it is drastically underestimated by the HH model. The excellent fit of the TM model suggests that the discharge of annihilating APs is predominantly driven by the rapid depolarization at the onset of AP generation. The failure of HH is in line with observations made in various studies, indicating that the rapidity of membrane dynamics is not well described by the HH model (***Cole and Moore, 1960; Baranauskas and Martina, 2006; Naundorf et al., 2006***). Our quantitative physical description of ephaptic coupling between neurons provides a theoretical framework for investigating biological systems in which the phenomenon of ephaptic coupling already has been described and to explore other neural connections. The general cable equation, linking field inhomogeneity to membrane current, applies to both source and target cell. This leads to a coupling between source and target, driven by the electric field and its inhomogeneities. The cable model directly explains why AP-induced membrane current in a neuron induces membrane current in targets (***Durand, 2014; Merrill et al., 2005; Tung, 2021***). Surprisingly, this straight forward concept has never been applied to ephaptic coupling at synapses. Virtually all numerical computations of ephaptic coupling rely on the HH model which, as shown above, is not suitable to predict the effects caused by AP annihilation.

Electrical fields of propagating APs create positive and negative currents in equal amounts and a propagating AP first hyperpolarizes, then depolarizes a parallel aligned target neuron. This may cause synchronization of APs and our proposed model also can be used to study the observed phenomena of synchronization due to ephaptic coupling, even in the case of zero discharge (see ***Figure 4***A, and local impact on the target, integrated on timescales >1 ms in ***Figure 5***A).

In contrast, AP annihilation in the source neuron acts as a strong local current injection. The non-zero discharge causes a more pronounced ephaptic coupling effect. AP annihilation occurs at the axon terminal and we have investigated ephaptic coupling for two neural connections to a greater extent. Both, the pre- and postsynaptic neurons have similar morphology and orientation and in addition, accessory structures at the site of connection. In teleost fish, an interneuron with an axon cap can have inhibitory, ephaptic coupling on the Mauthner cell’s axon initial segment (AIS) where APs are initiated (***Korn and Faber, 1975, 2005***).

In the vertebrate cerebellum, the Basket cells show ephaptic coupling with the Purkinje cell, and, comparably to the Mauthner cell, the generation of APs at the AIS is inhibited by a fast gating mechanism. In the following, we use the Basket cell – Purkinje cell connection as reference and for comparison to our model(s). As in all spiking neurons, an AP in the Basket cell generates a positive extracellular potential, and it is well documented that the Purkinje cell is very sensitive to extracellular potentials, since already 200 µV modulates its firing rate (***Blot and Barbour, 2014***). The cap structure (pinceau or basket (***Ramón y Cajal, 1909***)) around the terminal of the Basket cell restricts local currents to the Purkinje cell, increasing its sensitivity at the AIS to extracellular potentials (***Blot and Barbour, 2014***). The TM model predicts a fast and strong inhibition when APs are annihilating at the Basket cell (***Figure 4***B). Our numerical simulation revealed that in addition to the inhibitory effect of an annihilating AP at onset, a delayed (0.5 ms to 1.5 ms later) depolarization can occur at the target neuron. The magnitude of this depolarization depends on the time course of AP repolarization at the source (***Figure 4***C, ***Figure 5***). Presynaptic variation of AP width is a widespread mechanism for modulation of synaptic transmission by changing transmitter release, and ephaptic coupling possibly contributes to a direct modulation at the postsynaptic neuron (***Begum et al., 2016; Southan and Robertson, 1998; Kole et al., 2015***). ***Blot and Barbour (2014***) described a biphasic modulation of Purkinje cells with delayed and synchronized APs in Purkinje cells after ephaptic inhibition by the Basket cells. In their study, the drug GABAzine was used to investigate the controversially discussed role of GABA for this modulation (***Iwakura et al., 2012***). However, a biphasic modulation is consistent with the purely electrical framework provided in our RTM model, and it remains to be investigated whether presynaptic modulation of AP width is the underlying mechanism for modulating the biphasic ephaptic coupling of Purkinje cells.

Our finding and formal description of the strong ephaptic coupling generated by annihilating APs impose the need and provides the possibility to examine bioelectric effects in other areas. Endogenous electric fields can influence molecular processes within cells, leading to cell growth, maturation, migration and regeneration(***McCaig et al., 2000; Levin et al., 2017; Funk, 2015; Lyckman and Bittner, 1992***). The orientation of molecules and resulting structures can be induced by homogeneous electric fields and dielectric molecules can accumulate infield inhomogenities by dielectrophoresis (***Cifra, 2012; Pokorný, 2001***). In principle such effects can lead to persistent structural changes in neurons and thus may contribute to neural plasticity and memory. During synapse formation, presynaptic neurons interact with spines of the prospective postsynapse. Prior, or in parallel to chemical communication between the neurons that subsequently will form a synapse, Hebb’s rule might be implemented by a discharge from the source and coincident activity in the target. In this case, ephaptic coupling might be instructive for synapse formation. We included calculations on ephaptic coupling with the geometry of an end-to-end synapse and boutons on source/target (***Figure 4*** C-E) and we find that there is an initial sharp depolarization, followed by a slight hyperpolarization. Ephaptic coupling is highly amplified when source and target neurons have boutons, and such a morphology is omnipresent in spines and also presynaptic terminals commonly have enlargements as well. Our calculation further makes clear predictions where at the target (spine) depolarization can be expected, and this is the case about 1 µm away from the very tip of the target. It is important to highlight the importance of source–target geometry, predicted by our TM/RTM models, with opposite effects of ephaptic coupling in the two configurations: end-to-shaft and end-to-end.

Besides the annihilation of APs at the axon terminal, bidirectional propagation and hence collisions of APs might be more common than previously assumed (***Mateus et al., 2021; Scott et al., 2007; Debanne et al., 2011***). In theory, collisions can be used to perform computations (***Siccardi et al., 2016***) and neural networks possibly also perform such kind of information processing. For example, neurons for sound source localization perform a timing analysis based on binaural input. A neural network described as Jeffress delay line can accomplish such computation (***Jeffress, 1948***). ***Franken et al. (2021***) performed a delay sweep experiment and their findings on integration at MSO neurons (Medial Superior Olive neurons in vertebrates) look intriguingly similar to our collision experiment. Time differences and coincidence, in this case, is mapped on a location, e.g. the point of collision where a discharge with a center-surround profile is generated (***Figure 3***D) (***Treue, 2014***).

Irrespective of such speculative functionalities, ephaptic coupling is ubiquitous. Its effects span spectacular length and time scales, in some cases it can bridge up to hundreds of microns (***Kriebel, 1968; Chiang et al., 2019; Shivacharan et al., 2019***). Our formal description of ephaptic coupling between neurons provides a framework to study the functional significance of electric fields as a general mechanism for information processing in neural networks.

## Methods and Materials

### Experimental design

The objectives of the study were (i) to measure the electric field around propagating and colliding APs with unprecedented accuracy; (ii) to benchmark a powerful yet simple model of APs; (iii) apply this model to demonstrate its predictive power for ephaptic coupling in general.

### Sample preparation

The experiments were performed with the ventral nerve cord (VNC) of earthworms (*Lumbricus terrestris*). The specimen is placed in anesthesia (0.2% butanol in tap water) for 20 minutes. Then it is pinned in a basin, ventral side facing upwards, and covered with preparation saline (0.04% butanol in saline, 26 mM NaSO_4_ 25 mM NaCl, 6 mM CaCl_2_, 4 mM KCl, 1 mM MgCl_2_, 55 mM sucrose, 2 mM TRIS, adjusted to pH 7.4 (***Drewes and Pax, 1974***). The dissection starts with a small lateral incision caudally of the clitellum and it is followed by two longitudinal cuts alongside the VNC down to the posterior end. Afterwards the middle lappet is removed to lay open about 10 cm of the VNC. Gently pulling up the nerve cord reveals lateral connections which are then cut to disconnect the VNC from the rest of the nervous system. Once completely disconnected, the VNC is placed in chilled saline and kept at 4 °C for about 1 h.

### Electrophysiological recording of colliding APs

We use a custom made nerve chamber made from Polyoxymethylene which is encapsulated in a temperature controlled aluminum case (cover not shown, ***Figure 1***B) and kept at 12 °C. It contains a row of silverchloride electrodes (diameter 0.8 mm), located 8 mm above the bottom of the chamber and separated by 5 mm from each other. The recording chamber is pre-filled with chilled saline. The isolated VNC is transferred to the chamber and the saline is drained, leaving the VNC resting on the electrodes. Finally, the chamber is sealed with a plastic sheet and the aluminum case is closed. Two stimulators (Grass SD-9) are connected to electrodes at the ends of the nerve. In between, three custom amplifiers connect the recording electrodes to a digital storage oscilloscope (LeCroy WaveRunner 6050). Triggering of stimulation and recording is done with an Arduino micro controller. Data is acquired with 1 MHz, smoothed with a savitzki-golay filter (width 51 µs) and baseline corrected with the asymmetric least squares method (***Eilers and Boelens, 2005***). The VNC contains two giant fibers, the median (MGF) and lateral giant fiber (LGF). These are unambiguously distinguished by their individual stimulation threshold and propagation velocity. For the analysis we used 3 collision sweep experiments. Each experiment consists of numerous recordings from 3 channels and varying delay, the complete data of experiment 3 is shown in ***Figure 1—figure Supplement 1***.

### Models

#### General cable model for ephaptic coupling

The connection between inner electric potential ***V*** and transmembrane current is given by the general cable equation(***McNeal, 1976; Rubinstein and Spelman, 1988; Rattay, 1999; Anastassiou et al., 2010***).

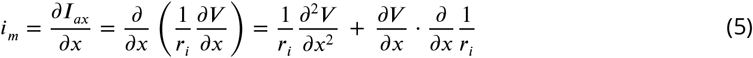

This equation describes both the generation of membrane current caused by an AP, and concomitant, the current induced by an external potential, as used in clinical applications (***Durand, 2014; Merrill et al., 2005; Tung, 2021***). In an infinite homogeneous neuron the membrane current is determined by the second derivative of the potential. The second term accounts for the spatial change of resistivity at any structural inhomogeneities, varicosities or neuron endings (***Basser and Roth, 2000; Holt and Koch, 1999***). Assuming a constant capacity and neglecting externalfields and inhomogeneities in the neuron one obtains the cable equation, eq. 1.

We assume the neurons being embedded in a large, homogeneous and isotropic conductor. In this case, the extracellular potential at a point *r* that is generated by the source neuron is given by

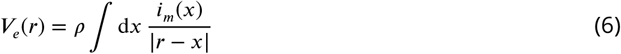

where *ρ* is the resistivity of the extracellular medium. The external potential ***V***_*e*_(*x, t*) adds to the membrane potential ***V*** _*m*_(*x, t*) of the target neuron via ***V*** = ***V*** _*m*_ + ***V*** _*e*_. Note that the target is considered to be in the resting state, where the transmembrane conductivity is negligible. The response is a redistribution of internal charges. In a static potential, the target neuron approaches a steady state by mirroring the external potential inhomogeneity in its transmembrane potential ***V*** _*m*_(*x*).

#### Relaxing Tasaki Model (RTM)

The repolarization of the extracellular potential missing in the TM model is added ad-hock by a repolarization function as follows. We introduce a state parameter *n*, which is 1 for the resting state. When the membrane voltage crosses the threshold value, *n* is set to 0 and its subsequent dynamic is given by

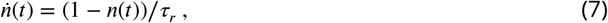

where *τ*_*r*_ is the repolarization time. The membrane parameters are controlled by

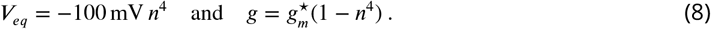

For very large *τ*_*r*_ (*≫ λ*^⋆^*/v*_*p*_), the RTM model reproduces the TM model, but eventually the resting state is restored a long time after an AP. With decreasing *τ*_*r*_ the repolarization affects the extracellular current and also the process of propagation. The exponent of *n*^4^ effectively causes a delay of the repolarization. The values *g* and *v*_*e*_*q* remain closer to the excited values for *t*^′^ < *τ*_*r*_*/*2, where *t*^′^ is the time since excitation. Under this condition the influence of the repolarization upon the initial rising phase is negligible. Consequently, the TM expressions for *v*_*p*_ and *λ*^⋆^ given above are in good agreement with the extended model (RTM).

#### Hodgkin Huxley (HH) model

The celebrated Hodgkin Huxley model also relies on the cable equation but adds a couple of additional equations to mimic the spatio-temporal shape of an AP (***Hodgkin and Huxley, 1952***). In particular, it incorporates the Nernst equilibrium and specific voltage and time dependent ionic conductances, resulting in a large number (typically of order 20!) of free parameters. This provides an enormous flexibility to account for almost any shape of AP. Nevertheless, our study reveals that the TM and RTM models which do not incorporate ion specific conductances are capable tofit experimental APs very well with only three degrees of freedom. We compare our data with the HH model for reference because of its widespread use and popularity. The complete equations and parameters are provided in ***Appendix 2***.

### Simulations

#### Fitting to experiment

For simulations of the APs we use the three models introduced above (TM, RTM, HH), all with the same general parameters and morphology. The python module BRIAN (***Stimberg et al., 2019***) is used to simulate the nerve with a multi compartment model using a forth-order Runge-Kutta method with time steps of 0.1 µs. After simulating the AP, the extracellular potential at the electrodes is calculated. In the nerve chamber, the nerve cord is hanging free between electrodes. The measuring electrode is surrounded by two ground electrodes which are 5 mm apart. All sections of the VNC that are in between the surrounding ground electrodes contribute to the measured potential according to

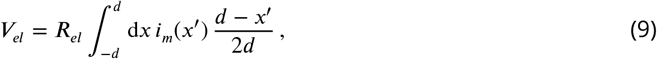

where *x*^′^ is the distance to the point of contact with the recording electrode, *d* is the length of the freely hanging nerve between two neighboring electrodes and ***R***_*el*_ is the resistance of the nerve cord between electrodes. We estimate the touching section of the nerve to be 0.8 mm (here *x*^′^ = 0) and, accounting for the sagging of the nerve between electrodes, we integrate over *d* = 5.2 mm in our calculations. In our approach, the magnitude of the TM model is controlled by 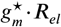. For easier comparison, we first adjust the TM model to the magnitude of the HH model, which is achieved with 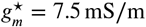. All parameters are presented in ***Appendix 3 — Table 2***.

Velocities are calculated from the time of AP arrival with an effective distance between the recording sites of 12 mm. The width of the collision is measured by the full width at half maximum of the negative deflection of the extracellular potential, as shown in ***Figure 1—figure Supplement 2***. The detailed settings as well as the calculated velocities and width are presented in ***Appendix 3 — Table 1***.

#### Predicted ephaptic coupling at synapses

We extended the standard BRIAN library to calculate the effect of an external field upon the target neuron by implementing eq. 5. The complete data for the end-shaft geometry with- and without bouton for all models is shown in ***Figure 4—figure Supplement 1***. Complete data for the end-end synapse is shown in ***Figure 4—figure Supplement 2***. We provide the full source code at osf.io/duyn3.

## Acknowledgments

We thank Mahlon Kriebel, Georg Raiser, Sabine Kreissl, Shamit Shrivastava, Gerardo Alvarez for fruitful discussions. This work was supported by the Deutsche Forschungsgemeinschaft (DFG) in the frame of the R. Koselleck project Ma 817/9 as well as from the Zukunftskolleg Konstanz.

## Data and materials availability

Source code and data have been deposited at the Center for Open Science (https://osf.io/duyn3/) Please refer to the readme file for further instructions.

## Appendix 1 The specific conductivity between the Basket and the Purkinje cell

The Pinceau is ensheathing the axon of the Purkinje cell. ***Blot and Barbour (2014***) report a resistance of ***R***_*pinceau*_ = 300 kΩ from inside the Pinceau to the surrounding ground. We can estimate the specific conductance (*ρ* in Ω m) by assuming an effective geometry. The Pinceau area is reported to be 100 to 600 µm^2^ (***Zhou et al., 2020***), e.g. a surface of 200 µm^2^ corresponds to a sphere of radius 4 µm. For a spherical shell with outer radius *a* = 4 µm and inner radius *b* = 3.5 µm we find

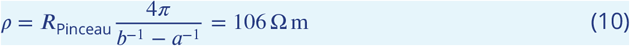

which is close to the 100 Ω m used by us. Please note that the numerical values used here are coarse estimates with large uncertainties, still the order of magnitude is well compatible with literature values.

## Appendix 2 HH model

The HH model (***Hodgkin and Huxley, 1952***) is implemented using standard parameters and expressions. We use the source code based on an example provided with the brian package (https://brian2.readthedocs.io/en/stable/examples/compartmental.hodgkin_huxley_1952.html, ***Stimberg et al. (2019***).) The total resistive current is given by

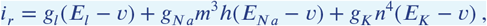

These three terms (channels) drive the membrane towards specific equilibrium values

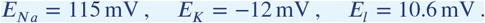

The conductivities *g*_*Na*_, *g*_*K*_, *g*_*l*_ are controlled by

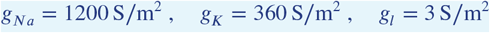

and

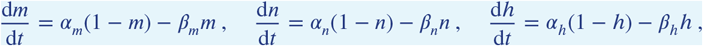

With

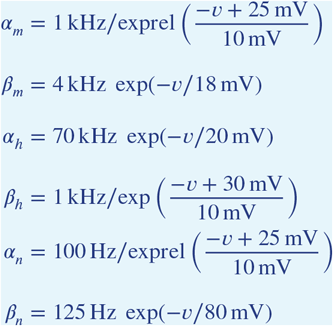

To avoid singularities, this implementation uses the function exprel which is provided by BRIAN (***Stimberg et al., 2019***), and is defined as

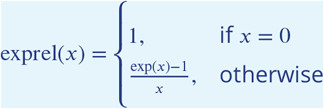

## Appendix 3 Additional experimental results

**Appendix 3—table 1.** Experiment details Settings as well as measured velocity and width of each experiment.

Additional experimental results
|  | exp. 1 | exp. 2 | exp. 3 | (unit) |
| --- | --- | --- | --- | --- |
| # recordings | 21 | 21 | 46 |  |
| delay range | 4 | 4.5 | 4.5 | ms |
| delay step | 0.2 | 0.23 | 0.1 | ms |
| sampling rate | 1 | 1 | 1 | MHz |
| duration | 10 | 10 | 10 | ms |
| $v_p$ C1→C2 | 16.2 | 16.2 | 16.3 | m/s |
| $v_p$ C1→C2 | 15.0 | 15.2 | 16.2 | m/s |
| $v_p$ C1←C2 | 15.1 | 15.1 | 14.9 | m/s |
| $v_p$ C1←C2 | 15.0 | 15.1 | 14.9 | m/s |
| $v_p$ C2→C3 | 13.1 | 13.1 | 13.3 | m/s |
| $v_p$ C2→C3 | 13.2 | 13.1 | 13.4 | m/s |
| $v_p$ C2←C3 | 13.0 | 14.3 | 14.5 | m/s |
| $v_p$ C2←C3 | 13.1 | 14.6 | 14.8 | m/s |
| width | 10.2 | 9.9 | 8.7 | mm |

**Appendix 3—table 2.**
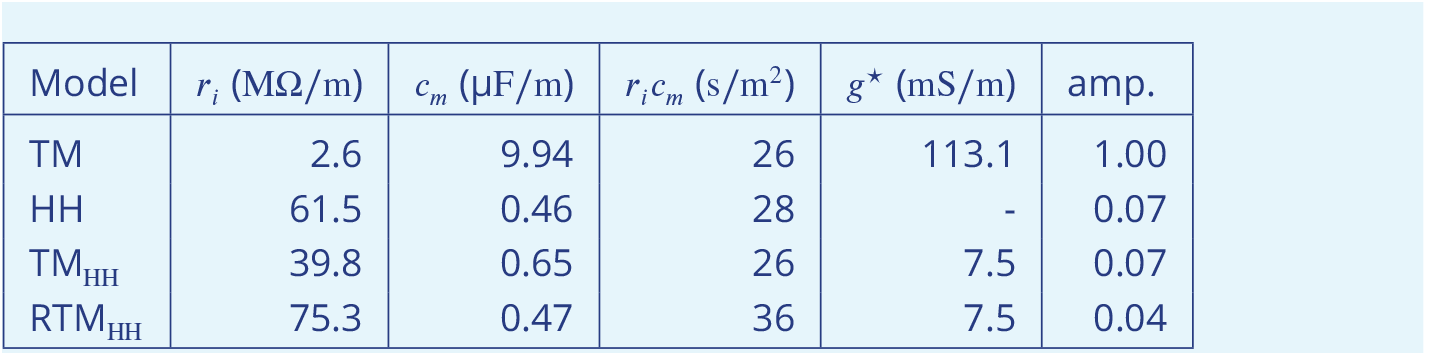
Model parameters For all models, we used the measured propagation velocity *v*_*p*_ (14.9 m/s) and the width of the collision, which is described by *λ*^⋆^ (1.8 mm) in the TM and RTM model, in order to adjust the parameters *r*_*i*_ and *c*_*m*_. For the HH model, we used literature values describing the different channels, their conductivities and time constants. For the TM model, we used the value of *g*^⋆^ as given by Tasaki and Matsumoto (2002). In order to compare the TM and the RTM with the HH model, we adjusted *g*^⋆^ such that the extracellular potential of the AP is comparable. Although the predicted amplitudes of the extracellular potentials are very different, the products *r*_*i*_*c*_*m*_ are very similar and in good agreement with literature values (see Tasaki and Matsumoto (2002)).

**Appendix 3—table 1. Experiment details** Settings as well as measured velocity and width of each experiment.
| Model | $r_i$ (MΩ/m) | $c_m$ (μF/m) | $r_i c_m$ (s/m<sup>2</sup>) | $g^*$ (mS/m) | amp. |
| --- | --- | --- | --- | --- | --- |
| TM | 2.6 | 9.94 | 26 | 113.1 | 1.00 |
| HH | 61.5 | 0.46 | 28 | - | 0.07 |
| TM<sub>HH</sub> | 39.8 | 0.65 | 26 | 7.5 | 0.07 |
| RTM<sub>HH</sub> | 75.3 | 0.47 | 36 | 7.5 | 0.04 |

**Figure 1—figure supplement 1.**
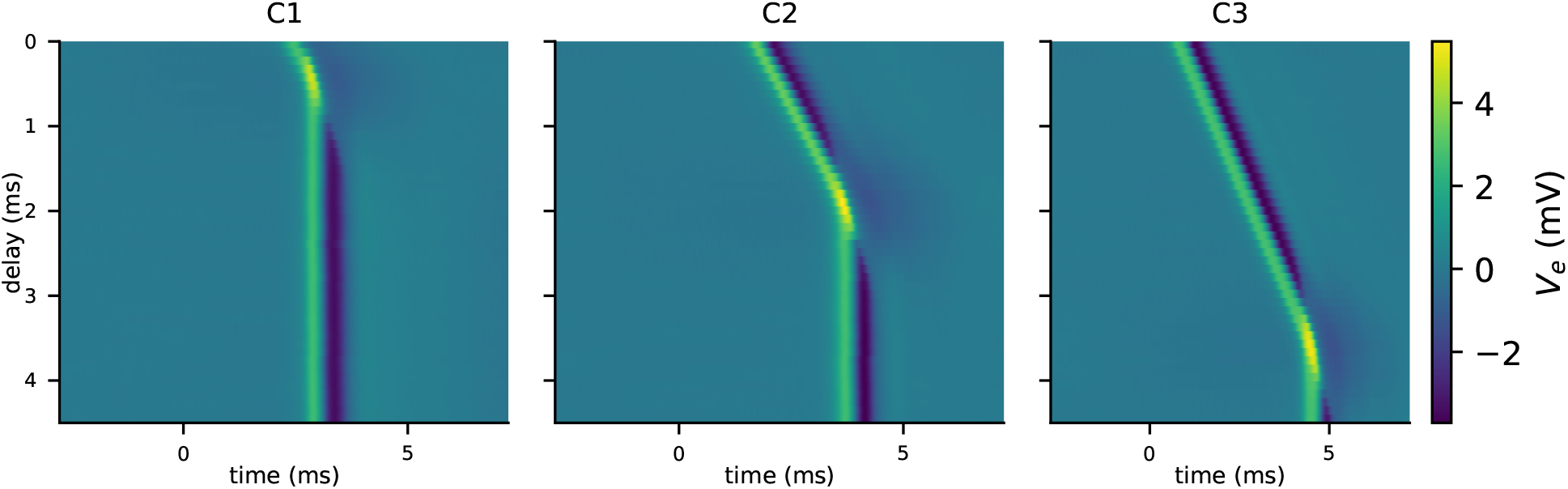
Complete raw data from experiment number 3 (out of 3).

**Figure 1—figure supplement 2.**
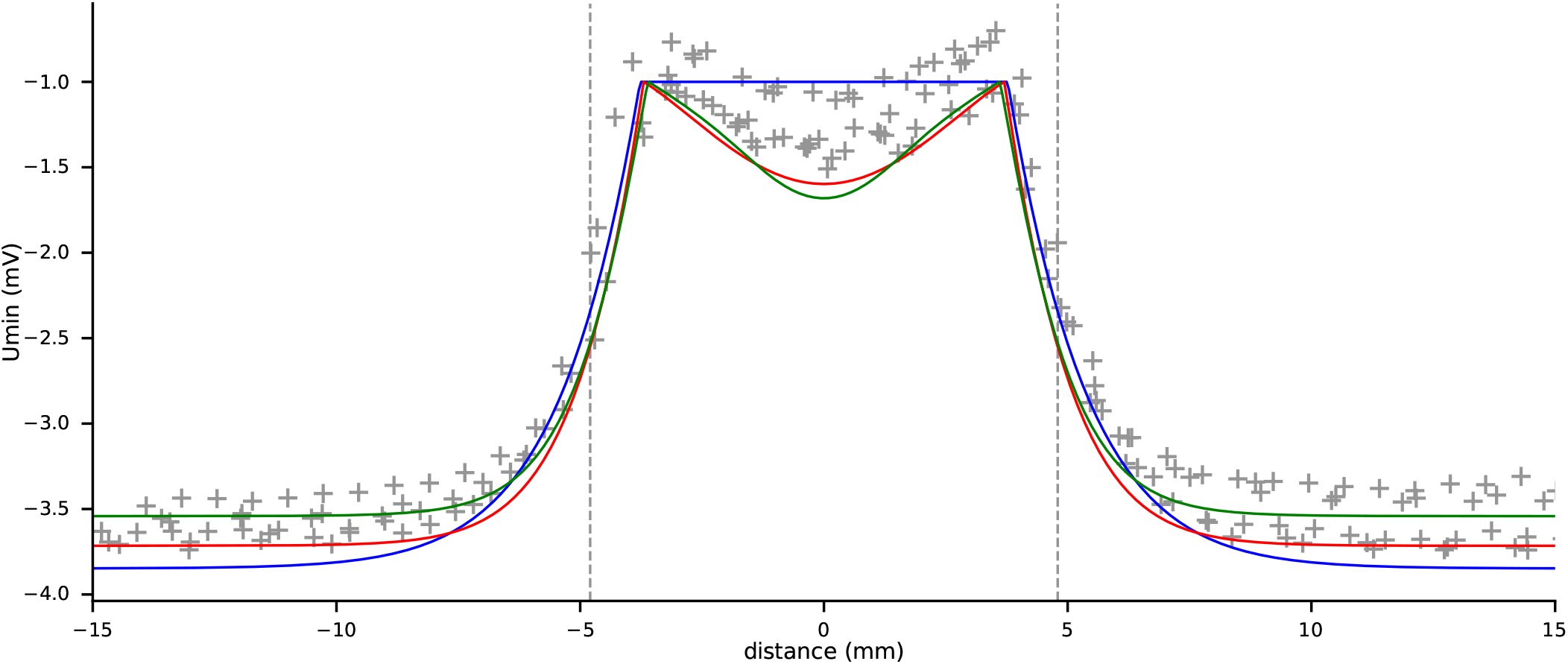
Grey crosses show the maximal negative deflection of the extra-cellular potential from 88 recordings (C1, C2, C3 from 3 experiments). Grey dashed lines denote the Full width at half maximum (from channel C2 from all 3 experiments), which is used to fit the models. Continuous lines are the TM model (blue), RTM model (red) and HH model (green). For better comparison, the simulated curves are shifted to peak at −1 mV.

**Figure 4—figure supplement 1.**
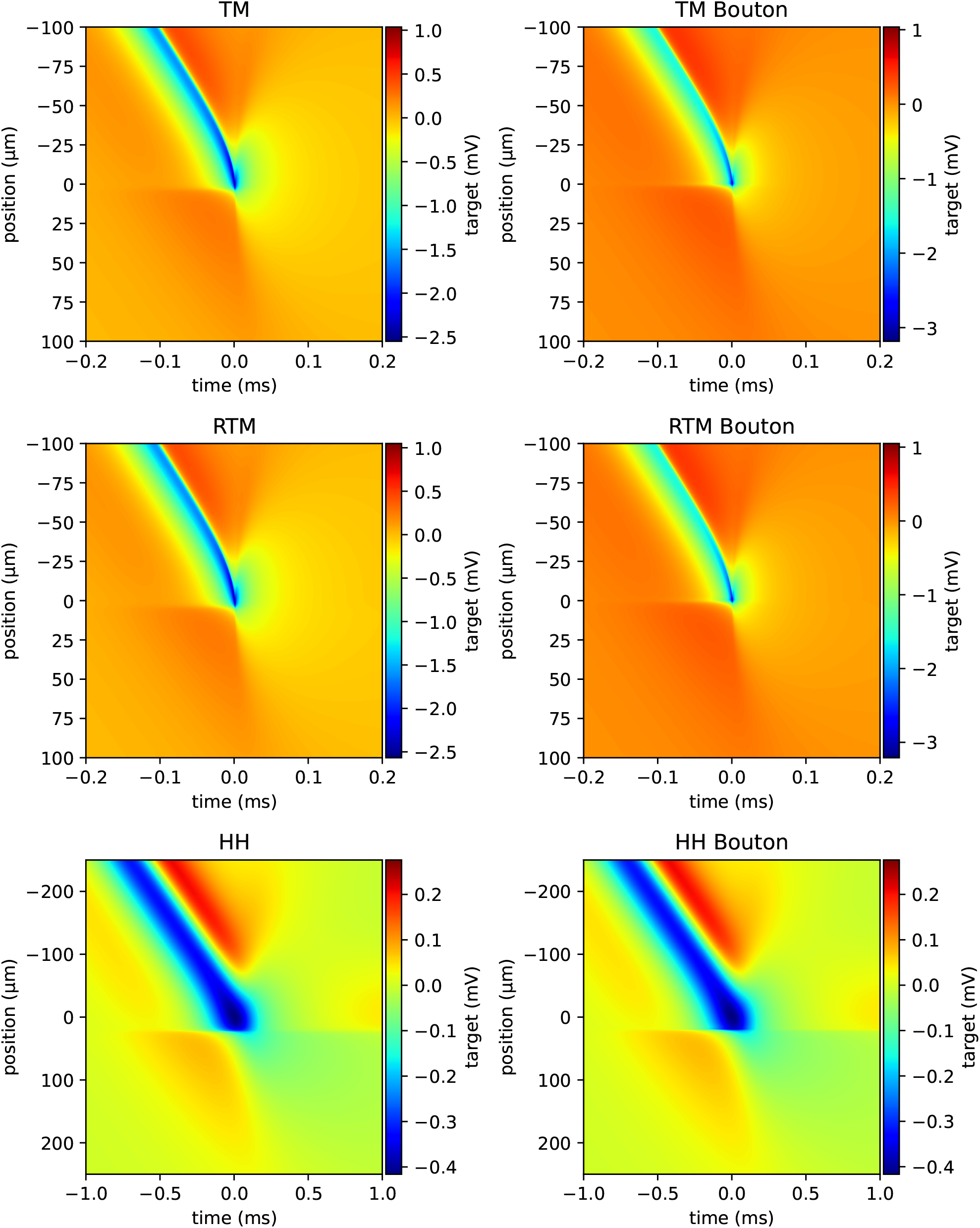
Simulation of target potential in an end-shaft synapse The y-axis denotes the position along the target where *y* = 0 refers to the target position next to the end of the source axon terminal.

**Figure 4—figure supplement 2.**
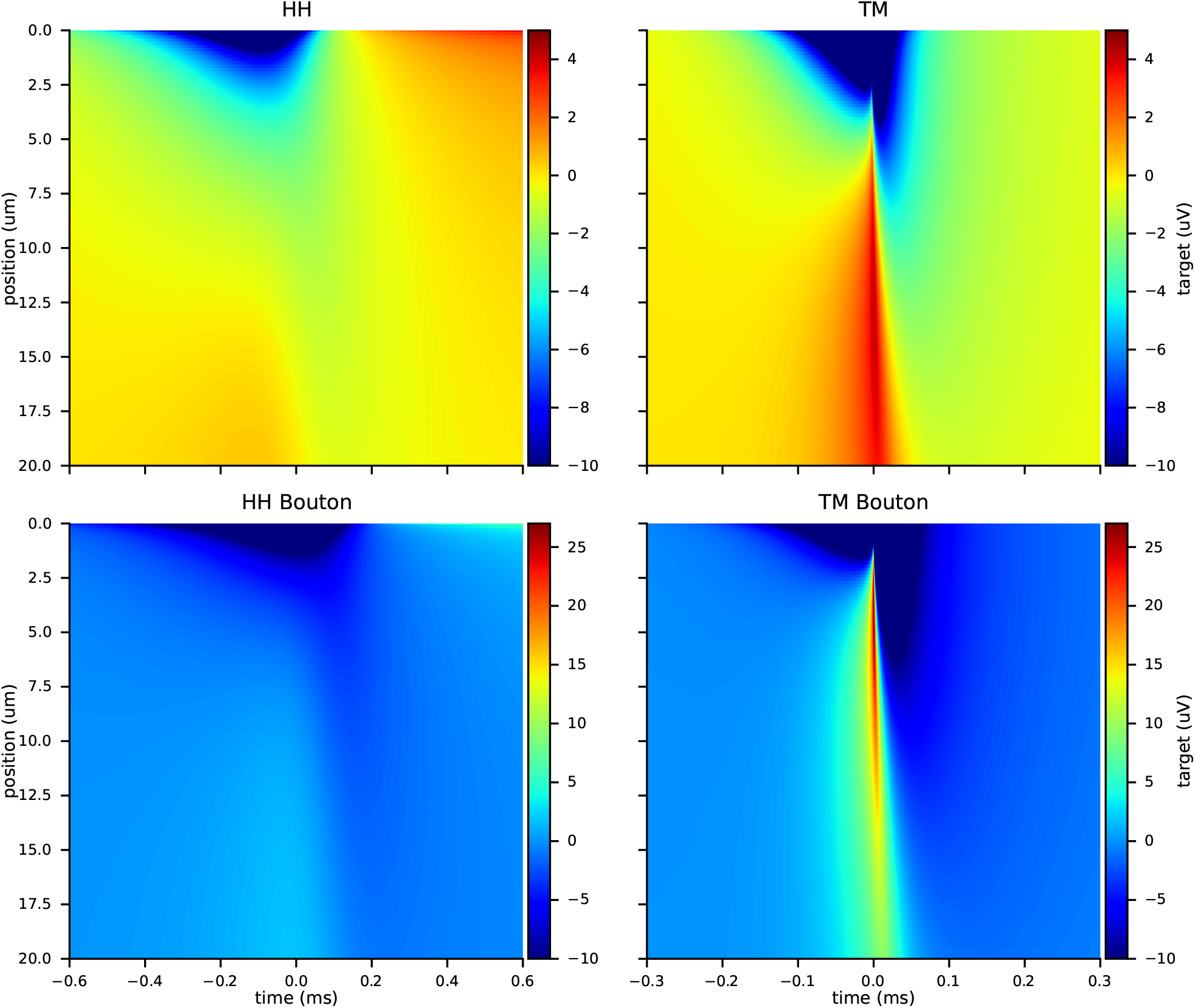
Simulation of target potential in an end-end synapse The y-axis denotes the position along the target where *y* = 0 refers to the tip of the target at the synaptic gap. The color scale is cropped at −10 µV.

## Notes

https://osf.io/duyn3/

### Competing Interest Statement

The authors have declared no competing interest.

### Summary of Updates

Added Fig 5A and corresponding sections in main text; Fixed a wrong unit in Appendix 3 -- Table 2

